# Molecular Counting with Localization Microscopy: A Bayesian estimate based on single fluorophore statistics

**DOI:** 10.1101/071191

**Authors:** D. Nino, N. Rafiei, Y. Wang, A. Zilman, J. N. Milstein

## Abstract

Super-resolved localization microscopy (SLM) has the potential to serve as an accurate, singlecell technique for counting the abundance of intracellular molecules. However, the stochastic blinking of single fluorophores can introduce large uncertainties into the final count. Here we provide a theoretical foundation for applying SLM to the problem of molecular counting based on the distribution of blinking events from a single fluorophore. We also show that by redundantly tagging single-molecules with multiple, blinking fluorophores, the accuracy of the technique can be enhanced by harnessing the central limit theorem. The coefficient of variation (CV) then, for the number of molecules *M* estimated from a given number of blinks *B*, scales like 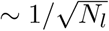, where *N_l_* is the mean number of labels on a target. As an example, we apply our theory to the challenging problem of quantifying the cell-to-cell variability of plasmid copy number in bacteria.

## I. INTRODUCTION

Cell biology is becoming increasingly quantitative with advances in light microscopy strongly driving this trend. Beyond imaging structure, significant effort has gone into developing microscopy based approaches to determining the abundance of proteins and nucleic acids in cells [1, 2]. Molecular counting experiments can yield additional insight into cellular structure and define the stoichiometry of interacting protein complexes. Moreover, since microscopy provides information at the single-cell level, it may be used to study stochastic variation within a population due to varying levels of mRNA and protein copy number, which is inaccessible to bulk techniques [3]. This variability is thought to be a crucial component of many biological processes such as cellular differentiation and evolutionary adaptation [4, 5].

A fluorescence based approach to molecular counting would be particularly powerful in single-cell ‘omics’ applications where a low level, such as trace amounts of protein, DNA, or RNA, must be detected [6, 7]. A reduction or even elimination of the amplification stage prior to sequencing of DNA or RNA could greatly increase the accuracy and reliability of single-cell genomic analyses. And since fluorescence microscopy is less susceptible to errors arising from protein size or abundance than techniques like mass spectroscopy [8], it could hold a significant advantage for single-cell proteomics.

Most conventional microscopy techniques either rely upon observing the step-wise photo-bleaching of fluorescent labels or on calibrating the fluorescence intensity to a standard [1, 2, 9]. Although these two methods have provided valuable insight into a range of cellular phenomena, both have their limitations. Step-wise photobleaching can only be used to identify small numbers of molecules (roughly < 10). And intensity measurements, although able to quantify the number of more abundant molecules, are hindered by stochastic variation in photon emission and collection efficiency, and are limited by the dynamic range of the detection camera. Likewise, both techniques have difficulties when observing diffraction limited fine structures due to overlapping signal from neighbouring features.

Super-resolved localization microscopy (SLM), which include techniques such as PALM [10] and dSTORM [11], could provide an alternative approach that would not suffer from these limitations. SLM can produce images of structural detail an order-of-magnitude finer than diffraction limited techniques. The method relies on precisely localizing the spatial position of single, fluorescent labels attached to an assembly of target molecules. This typically requires the use of photo-convertible or photo-activatable fluorophores that can be induced to blink in such a way that only a random subset of the labels are visible during each frame [12, 13]. For a sufficiently sparse image, each diffraction limited spot should be sufficiently well separated, and the subset of fluorophores may be localized with a precision that scales like 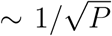, where *P* is the mean number of photons collected from a single blink of a fluorophore. Tens of thousands of frames are typically acquired, the spatial coordinates of the fluorophores within each frame extracted, and the resulting data from the stack rendered into a final image.

Since SLM measures discrete blinks from single fluorescent labels, it essentially provides a digital approach to molecular counting, compared to conventional techniques that measure the overall amplitude of a signal, and are akin to an analog method [14–16]. By focusing on interpreting the number of detected blinks, the usefulness of SLM moves well beyond what can be achieved with imaging alone. For instance, intracellular elements like multimerized membrane bound proteins, which are still unresolvable by SLM imaging, could be detected. Likewise, this approach relaxes the spatial accuracy requirements of imaging, opening the way for faster detection, at lower signal, and on smaller detector pixel arrays. However, there are several challenges to obtaining accurate counts with SLM, most notably, accounting for multiple blinks from a single fluorophore and the inefficiency with which the fluorophores photo-activate or photo-convert [17, 18]. Both issues lead to an inaccuracy in estimating the total number of molecules [15, 16, 19], and there has been much effort to mitigate these difficulties [14–16, 20–25].

Starting from the statistics of the observed number of blinks of a single fluorophore, our approach is to apply Bayesian analysis to estimate the number of molecules from the total number of blinks detected in an SLM measurement (a related, but distinct approach, is presented in [26]). We are able to derive an analytic expression both for the estimated number of molecules and the error in that estimate. In addition, although the stochastic blinking of the fluorophores can introduce uncertainty when translating between the number of localizations and the number of molecules, we show that labeling single-molecules with multiple labels can reduce this uncertainty. As an example, we apply our theory to design an experiment that measures the cell-to-cell variability of plasmid copy number in bacteria (a task that has proven to be surprisingly difficult [27]).

## II. METHODS

### A. Counting single molecules from blinking fluorophores

Let’s begin by calculating the conditional probability distribution *p*(*B*|*N*) for observing *B* blinks (or localizations) from a set of *N* fluorophores during a measurement time *T_M_*. We assume the only information available is the total number of blinks *B*, and ignore any spatial information contained within the data that might enable us to differentiate one fluorophore from another. In the simple case of a single emitter, the probability *p*(*B*|*N*) is often well approximated by a geometric distribution:

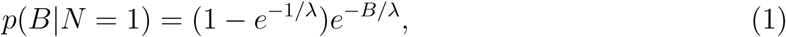
where *λ* is the characteristic number of blinks of a particular fluorophore within the interval *T_M_*. This distribution arises when the blinking is a Poisson process between an ‘on’ and an ‘off’ state that after sufficient time is truncated by photobleaching. From this relationship we generalize to the case of *N* fluorophores to obtain a negative binomial distribution

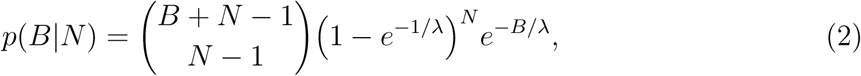
where the prefactor accounts for the number of ways that *N* fluorophores, each blinking some B*_i_* times, can yield 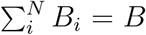 blinks. The mean and variance of Eq. 2 (see Appendix A) are:

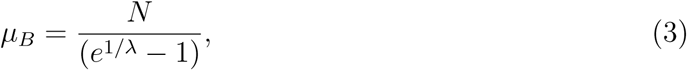
and

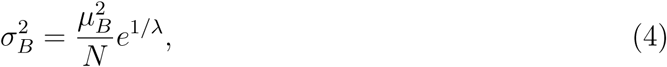
respectively.

Up until this point, we have been considering the conditional probability distribution *p*(*B*|*N*), which, to reiterate, is the probability of observing *B* blinks when there are *N* fluorophores. However, we wish to know the probability of there being *N* fluorophores when we observe *B* blinks, or *p*(*N*|*B*). In the language of Bayesian statistics, we need to connect the likelihood *p*(*B*|*N*) to the posterior distribution *p*(*N*|*B*), which can be achieved by Bayes’ theorem:

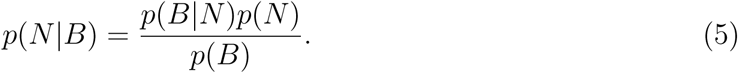
If we have no prior knowledge of the distribution of fluorophores in our sample we may set the prior *p*(*N*) as a constant [28]. The posterior and the likelihood are then proportional *p*(*N*|*B*) ∝ *p*(*B*|*N*).

We define a log likelihood function 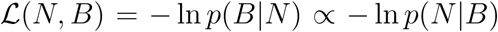 and apply a Laplace approximation to the posterior distribution. That is, for a sharply peaked, symmetric distribution, the maximum with respect to *N* (i.e.,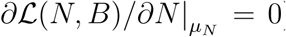) should roughly correspond to the mean number of fluorophores:

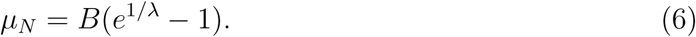

Likewise, we can obtain the variance in the estimated number of fluorophores, which provides the accuracy of the estimate, by rewriting the posterior distribution and Taylor expanding as follows:

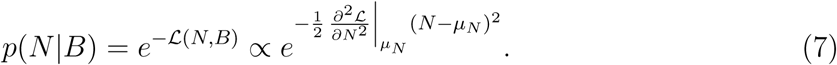

In the exponent of Eq. 7, we identify the estimator of the Fisher information matrix [29] 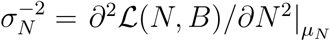 to yield the variance

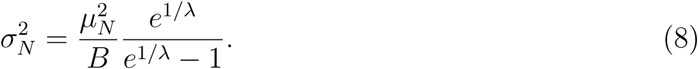

For fluorophores that blink multiple times during the measurement (i.e., the limit λ ≫ 1), Eq. 6 simply reduces to the intuitive expression μ*_N_* = *B/λ*, which states that the most likely number of fluorophores is equal to the measured number of blinks divided by the mean number of blinks per fluorophore. In this limit, Eqs. 6 and 8 approach the Poisson limit with variance 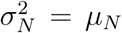, and the coefficient of variation (CV), which quantifies the variability of the estimate relative to the mean (*η* ≡ *σ_N_/μ_N_*), is simply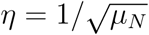.

### B. Accounting for multiple labels on a target

There are a myriad of labeling techniques in cell biology and the correspondence between the number of fluorophores and the number of target molecules is typically not one to one. For instance, immunolabeled molecules will contain several dyes on each antibody and covalently labeled proteins will often be tagged at multiple residues. The probability of having *N* fluorophore labels in total when there are *M* target molecules, each with *h* possible sites where a fluorophore may bind (or hybridize), is given by the binomial distribution

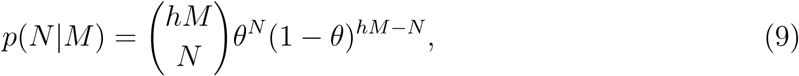
where *θ* denotes the fractional occupancy. Note that *hM* is the maximum number of labels possible, if we ignore all non-specific labeling, and that the fractional occupancy *θ* is always less than one.

### C. Distribution of blinks within a population

We can now combine Eqs. 2 and 9 as follows:

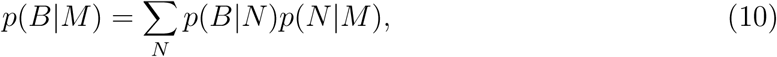
to derive the conditional probability distribution for observing *B* blinks from a population of *M* fluorescently labeled molecules. Although the full sum is quite cumbersome, if straightforward to evaluate numerically, the moments of Eq. 10 are analytically tractable. For instance, the first and second moments may be found by multiplying both sides of Eq. 10 by ∑*_B_ B* or ∑*_B_ B*^2^, respectively, and evaluating the summations. The first moment is the mean number of blinks

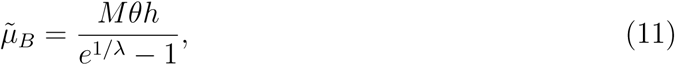
which can be combined with the second moment to obtain the variance

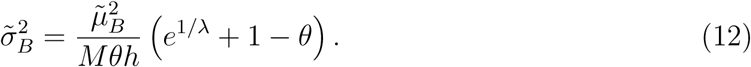

However, we wish to estimate the mean and variance in the estimate of the number of molecules after having measured *B* blinks. Although a more formal derivation is provided in Appendix B, the estimate for the mean can simply be obtained by substituting 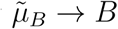 and 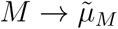 into Eq. 11:

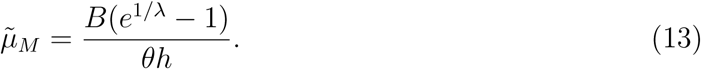

In the limit λ ≫ 1, Eq. 13 again yields an intuitive result for the expected number of molecules 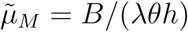.

The variance, on the other hand, is more challenging to evaluate, but it can be estimated, similar to how one estimates the propagation of errors in a measurement (see Appendix C).

If we assume the distribution *p*(*M|B*) is peaked about the mean 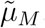, then the Fisher information matrix is:

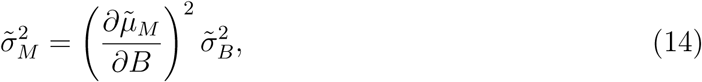
to yield our final result for the estimate of the variance

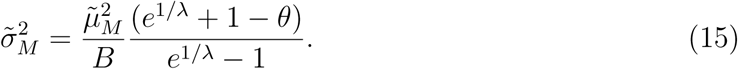

In the limit λ ≫ 1, this yields the simpler expression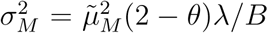, and the CV is simply

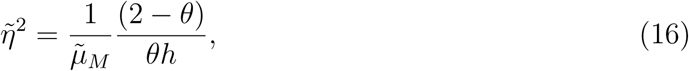
which can, when *h* > (2 − *θ*)/*θ*, reach the sub-Poissonian limit scaling like one over the square root of the mean number of labels per molecule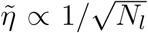, where *N_l_* ≡ *θh*. This scaling, of course, is simply a result of the central limit theorem.

## III. RESULTS

### A. Cell-to-cell variability of plasmid copy number

To illustrate the utility of our approach, we consider the problem of counting plasmids in single bacterial cells. Plasmids are circular, extra-chromosomal segments of DNA that often confer a selective advantage, such as a resistance to antibiotics, to their host. Plasmids are also a relatively simple way to introduce genes into a cell making them an invaluable tool throughout molecular and synthetic biology. An important feature of a plasmid is its copy number. If a plasmid is harbouring a gene one wishes to express at a controlled level, variations in plasmid number will likely lead to varying levels of expression. Although bulk techniques such as qPCR can place bounds on the average plasmid copy number, it has proven extremely difficult to quantify the copy number distribution within a population [27].

Localization microscopy could provide a way to measure the cell-to-cell variability in copy number. Super resolved localization microscopy images of a high-copy number ColE1 plasmid were recently obtained in fixed Esherichia coli bacteria [30]. Atto-532 labeled DNA probes were annealed via DNA fluorescence in situ hybridization (FISH) to an array of LacO sites (256 sites) introduced in the target plasmids, then imaged by dSTORM. Furthermore, both the mean number of blinks λ and the fractional occupancy *θ* could be obtained from *in vitro* measurements (from photoactivation of sparse samples of the dye and from photo-bleaching experiments on the hybridization of the probes to an array of the target sequence, respectively).

Here we consider targeting a 96 site array (a 96-*TetO* array, for instance, is commonly available [31]) to lessen the effects of the insert on the replication dynamics of the plasmid [32]. Figure 1 shows the probability distribution *p*(*M*|*B*) for this hypothetical system. We’ve chosen *M* = 25 for illustrative purposes, but the qualitative results remain similar for different plasmid number (i.e., for increasing fractional occupancies, *Nl/h* = θ, the distribution becomes increasingly peaked around the expected value). Figure 2 shows that as more blinks are observed, due to an increased fractional occupancy, for the same number of plasmids (*M* = 10, 25, 100), the error in the estimate of *M* rapidly decreases. As more probes associate with the plasmids, the coefficient of variation decreases like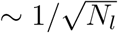, and can be made to drop well below the Poisson limit 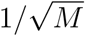(see Fig. 3). This is illustrated for a range of plasmid number (again, *M* = 10, 25, 100). For *M* = 25, for instance, and a reasonable fractional occupancy as might be achieved by DNA FISH, say 20%, the error in a single-cell count would be only ±1-2 plasmids. Unfortunately, the efficiency at which the probes hybridize in DNA FISH experiments is always hampered by the competing complementary DNA. Perhaps by employing peptide nucleic acid (PNA) probes [33], devoid of the negative charge along their backbone, the fractional occupancy could be enhanced to further reduce the uncertainty.

**FIG. 1.**
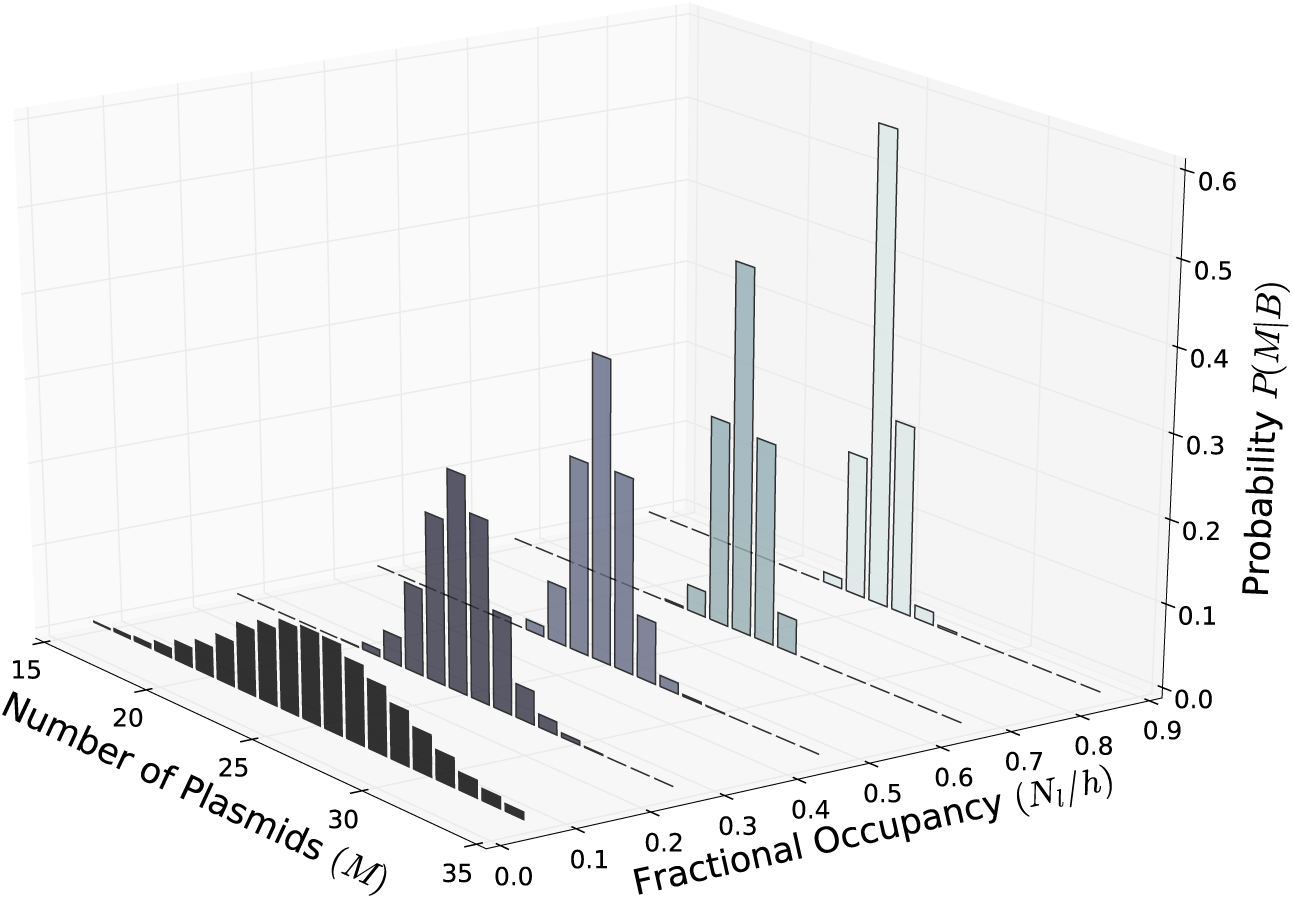
The simulated probability distribution *p*(*M*|*B*) for (*M* = 25, λ = 2, *h* = 96) at fractional occupancies (*N_l_/h* = 0.05, 0.25, 0.45, 0.65, 0.85) showing a sharper distribution (i.e., less uncertainty) at increasing values.

**FIG. 2.**
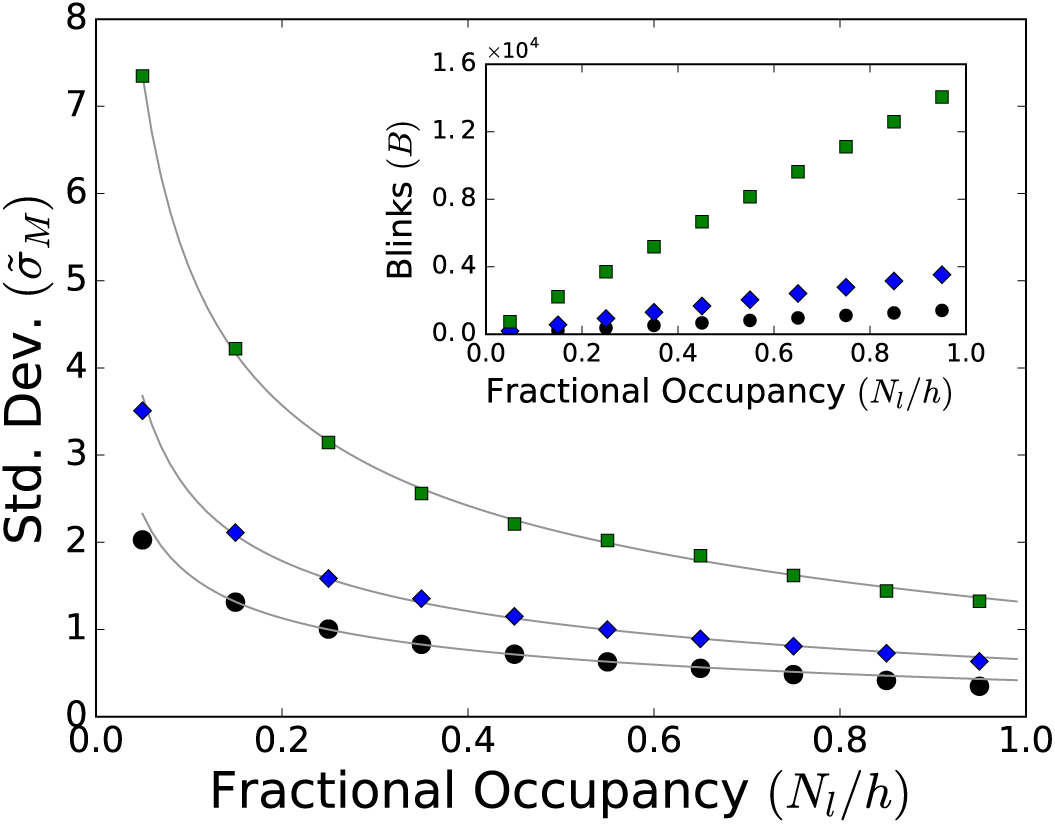
Uncertainty in the number of molecules vs. fractional occupancy for *M* = 10 (circles), 25 (diamonds), and 100 (squares), λ = 2 and *h* = 96. Solid lines are a theoretical estimate from Eq. 15. The insert shows the increase in the expected number of observed blinks, for these parameters, with increasing fractional occupancy.

**FIG. 3.**
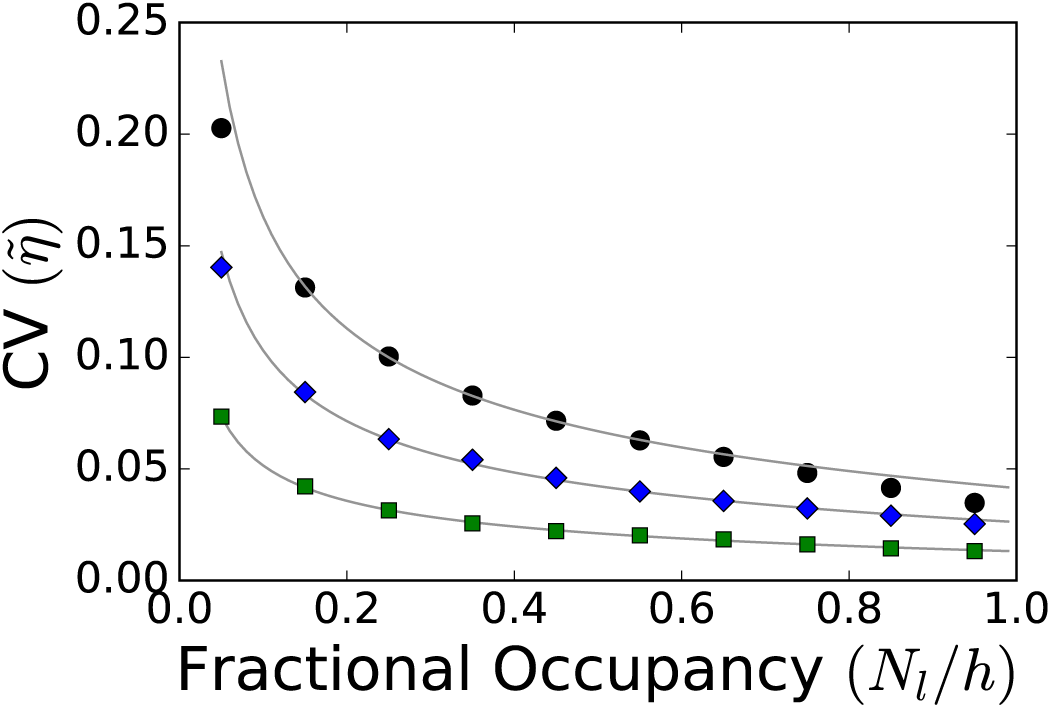
Coefficient of variation (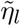) vs. fractional occupancy (*N_l_*/*h*) for different plasmid (molecule) number: *M* = 10 (circles), 25 (diamonds), 100 (squares) (λ = 2 and *h* = 96). The solid lines are the theoretical estimate from Eq. 16. At, roughly, *M* = 10 the theory begins to deviate from the simulated results.

### B. Realistic *In silico* single-molecule counting

In practice, a range of considerations must be accounted for before the theory we’ve developed above can be applied; the most important consideration being to generate an accurate table of single-molecule localizations. To avoid tackling all the complications of SLM counting concurrently, we first develop a practical approach to molecular counting on simulated realizations of an SLM counting experiment. For instance, our theory assumes that all the fluorophores blink according to a geometric distribution characterized by a single parameter λ, but if the sample is not homogeneously illuminated, or if the local chemical environment varies across a sample or with time, this criteria might not hold. Our *in silico* data allows us to impose temporal and spatial uniformity in the blink statistics. Likewise, *in silico* we know exactly how many fluorophores we are attempting to count and don’t have to calibrate for a sub-population that refuses to photoswitch (a complication our theory does not incorporate). All these issues can be avoided, for the moment, by analyzing simulated images. The images are then processed to generate a localization table, just as one would process actual SLM data, and from the resulting localization table we show how to extract molecular counts according to the theory. In what follows, we essentially model dSTORM data with an organic dye, and many of the parameters are taken from our experimental setup (e.g., pixel size, frame rate, etc.).

We model the switching photophysics of each fluorophore as a Poisson process truncated by a geometric distribution [23]. The Poisson process is an effective model of photo-switching between an ‘on’ and an ‘off’ state by the fluorophore, which is only able to switch a limited number of times, drawn from a geometric distribution, before photobleaching. We choose an ‘on’ time t_on_ = 100 ms and a duty cycle *t*_on_/*t*_off_ = 10^−3^, which specifies the Poisson dynamics. As to the bleaching dynamics, we set the characteristic number of blinks λ = 2, which is a reasonable value [30]. ‘On’ states are rendered with a Gaussian point spread function (PSF) of width σ_PSF_ = 127 nm and Poissonian, shot noise intensity fluctutations (mean 1750 photons/frame). Individual blinks are then pixelated by the finite size of the camera pixels (1 pixel =117 nm) and white noise is added to the images to account for background (mean = 48, std = 10, photons). Finally, the dynamics, evaluated at 1 ms time-steps, are discretized into 50 ms frames to provide a stack of *in silico* data for processing.

Despite knowing the exact value of λ (since it’s a parameter in the simulation), we first attempt to measure λ as one might in an actual experiment. We start with a grid of 1156 fluorophores each spaced 7 pixels apart, and the simulation is run for a total simulated time of 12.5 minutes, which yields 15,000 images in the stack. The images are then analyzed with RapidStorm (a popular, open-source package for localization microscopy [34]) using fixed-width Gaussian fits to the PSF. Once we have built an initial table of localizations, we then identify blinks that last for multiple frames as a single blink of a fluorophore by associating all temporally, consecutive localizations within a radius of 100 nm [35]. Since the initial grid of fluorophores is well separated, we can now build a histogram of the number of blinks from a single fluorophore and fit to obtain λ. An example of the resulting distributions is shown in the inset to Fig. 4.

**FIG. 4.**
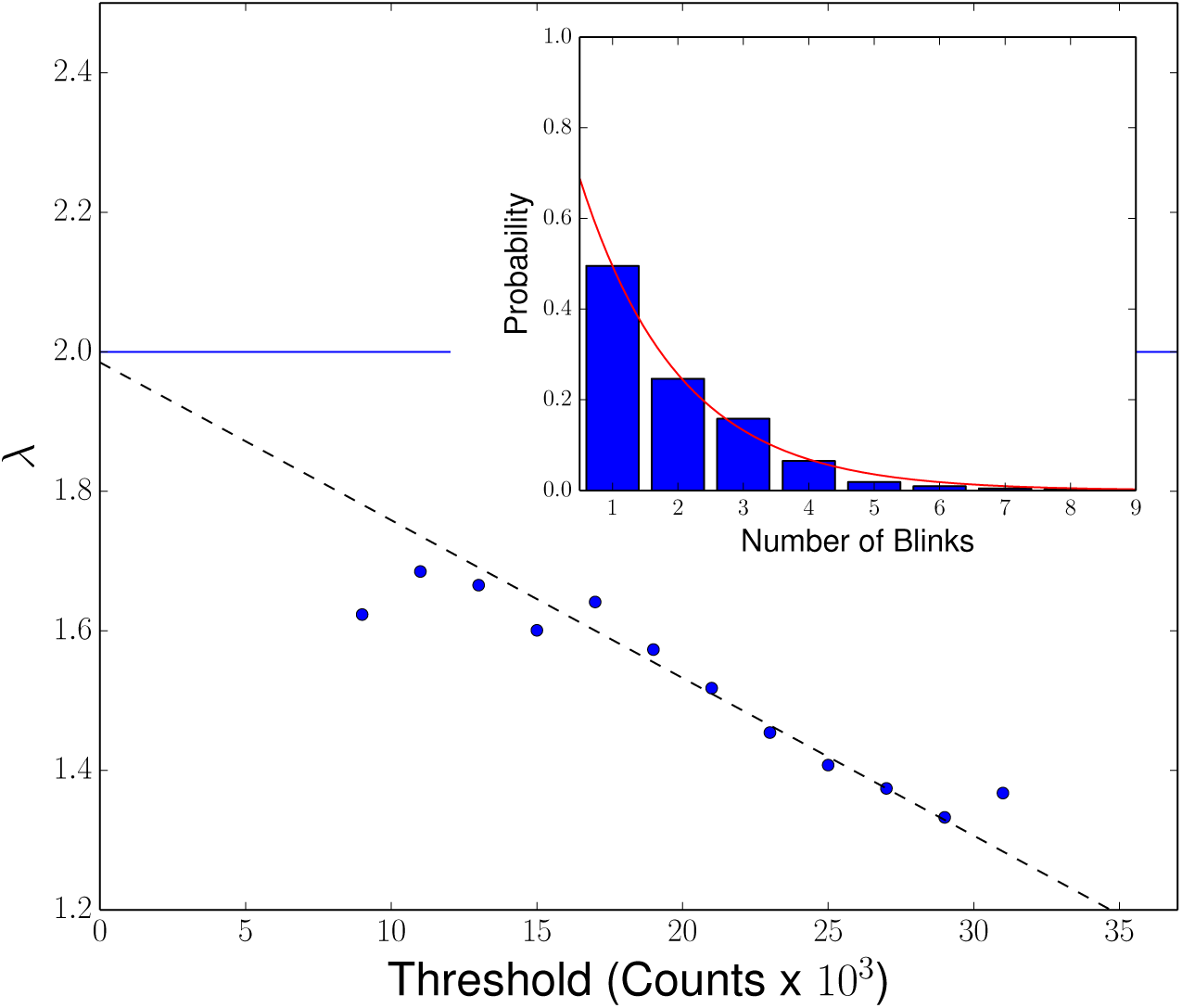
Plot of λ vs. Threshold. The dashed line is a linear fit to the data points from 15-25×10^3^ counts. The solid blue line is the simulated value λ = 2, which well agrees with the extrapolation of the measurements at zero threshold. The insert shows a typical histogram of the number of blinks from a single fluorophore (for this example, the threshold was 21×10^3^ counts). The red line is the geometric fit from which we extract λ.

Note, the total number of localizations is sensitive to the choice of threshold, which in RapidStorm is the integrated number of camera counts (here, 47 counts/photon) within the fitted PSF. For too low a threshold, it becomes difficult to differentiate a blink of the fluorophore from background noise. In fact, setting too low a threshold will generate a localization table with many spurious, random localizations. Our solution is to evaluate λ at increasing thresholds and extrapolate to the zero threshold value. That is, at different thresholds, we build a histogram of the number of blinks from a single fluorophore, fit to a geometric distribution to obtain λ, then plot λ vs. threshold. For a range of intermediate thresholds, λ steadily decreases as the threshold is increased. As shown in Fig. 4, linearly extrapolating the data to zero threshold gives excellent agreement with the value we explicitly coded into the simulation (1.98 compared to 2).

With this calibration in hand, we next turn our attention to the actual *in silico* molecular counting experiment. We build a grid of 1225 vertices, each 8 pixels apart, and scatter about each vertex (within a radius of 2 pixels) up to *h* = 4 fluorophores (the actual number chosen from a binomial distribution with fractional occupancy *θ* = 0.75). The procedure is similar to our calibration experiment: we simulate the system for 15,000 frames with the same parameters as before, analyze the stack in RapidStorm, bunch together blinks that last for multiple frames, and repeat at increasing threshold. To obtain the total number of blinks *B*, we determine the total number of blinks at each threshold, and again linearly extrapolate to zero threshold (see Fig. 5). This result can be used in Eq. 13 and Eq. 15 to obtain the mean number of molecules 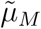 and the variance 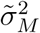 of our estimate. Alternatively, we could directly plot Eq. 13 and again extrapolate to zero threshold (see Fig. 5).

**FIG. 5.**
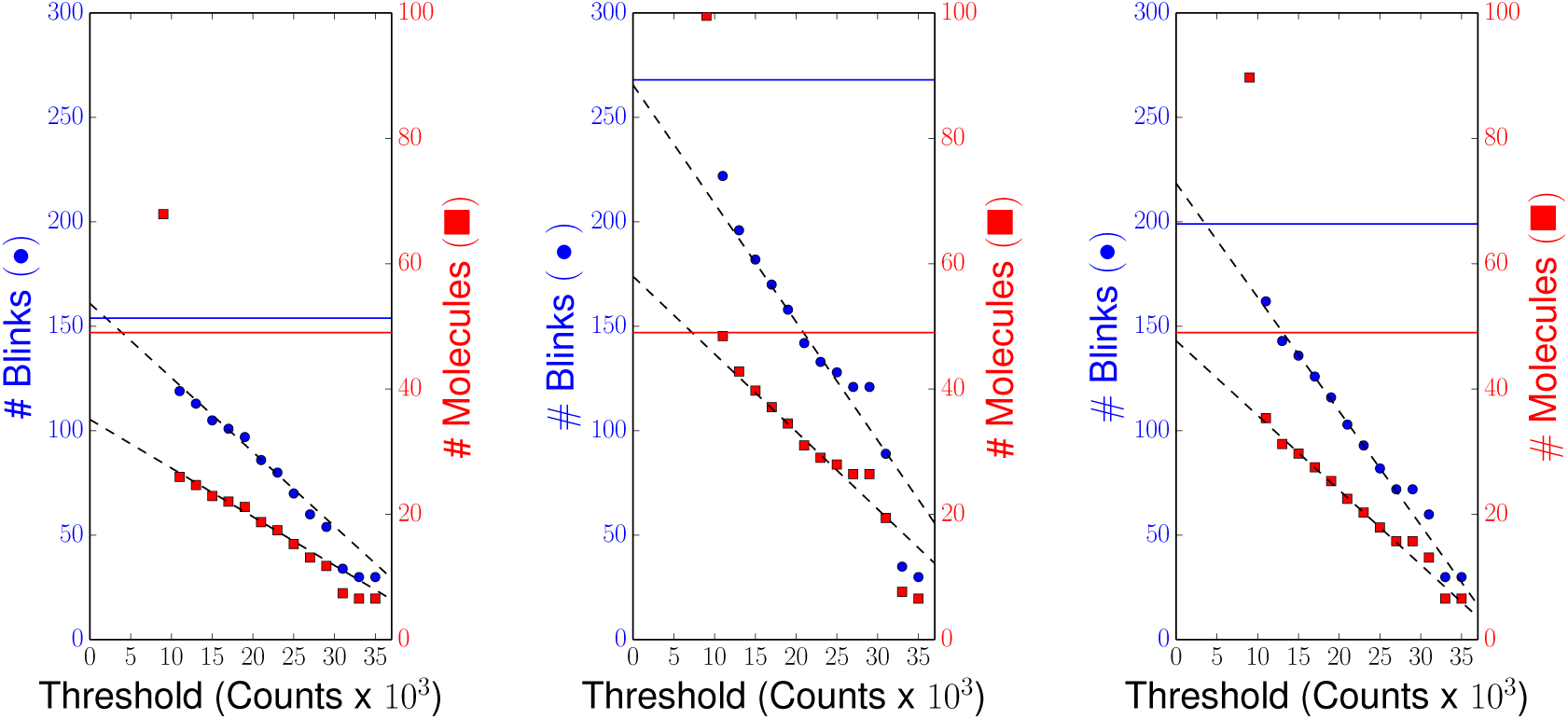
Total Number of Blinks (Molecules) vs. Threshold for three realizations of the *in silico* experiment. In each figure, the blue circles (red squares) are the measured number of blinks (molecules) at each value of the threshold. The solid blue (red) line are the actual number of blinks (molecules). Dashed lines are linear extrapolations of the data fit to a threshold range of 15-25×10^3^ counts.

From 25 realizations of this experiment, our measurements yielded 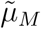 = 45±9 molecules with the actual number of molecules fixed at 49. The error on our estimate is slightly larger than our theoretical estimate from Eq. 15 (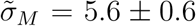), but this is largely due to the inherent error in extrapolating the data to zero threshold. Regardless, our estimate on the molecular count is still within 20%, which is not bad considering the stochastic nature of the system we’ve considered. Note, as the background noise is increased, it will become increasingly difficult to reliably extract the localizations and may make the approach we’ve just presented impractical. Likewise, the density of labels we consider in this section is still relatively sparse. In dense samples (or in samples where one might want to rapidly acquire the data by reducing the ‘off’ time), it may be hard to guarantee that the blinks don’t overlap within a frame, skewing our estimates of the total number of blinks. Improved software for localizing single-molecules, in noisy and/or dense samples, may help limit these artifacts [36, 37].

## IV. DISCUSSION

Our approach relies upon an accurate measure of two parameters: the mean number of blinks from a single fluorophore (λ) during the measurement time, and the fractional occupancy (*θ*). Although it is by no means certain, as a starting point, lets assume that *in vitro* measures of these parameters are accurate. The parameter λ can be obtained from imaging single blinking fluorophores sparsely attached to a coverslip. Of course, inherent in this measurement is the assumption that the blink statistics are both spatially uniform and temporally invariant across the sample (or, at least, the region of interest). It should be kept in mind that this is not always the case: flat-field illumination is a challenge, densely packed labels may interact through charge/energy transfer, there may be pH and other local environmental variations, and so on.

Obtaining the fractional occupancy is another challenge [38]. One method is to count photobleaching steps of single labeled probes bound to a target. In our plasmid example, the actual 96-*TetO* repeat target is much too large for such an approach, but it’s reasonable to assume that a smaller (say, 10-15 repeat target) would extrapolate. Although this would provide the full distribution of the occupancy, a simpler strategy is to measure the resulting ratio of plasmid DNA to fluorophore labels (by absorption spectroscopy and fluorometry, respectively). The fractional occupancy can then be backed out of the underlying binomial distribution. Of course, a binomial distribution is only an approximation to the occupancy statistics, and assumes that it’s equally likely for a labeled probe to associate with any one of the complementary binding sites along the plasmid. If the probes were to interact (e.g., electrostatic or steric interactions), for example, the underlying distribution may be more complicated than our simple model assumes.

Moreover, as mentioned, many fluorophores do not efficiently photo-activate or -convert. One might be be able to account for this aspect of the photophysics by quantifying the fractional occupancy using DNA origami [39]. For instance, in our plasmid example, an array of *TetO* sequences that could be spatially resolved by localization microscopy (e.g., patterned on a grid) would serve as a template. Inefficient photo-activation or -switching will simply lead to a reduced, measure of the fractional occupancy. Given a sufficient number of hybridization sites *h*, this should account for any underestimation in the molecular number due to inefficient photoswitching. In fact,an added advantage of labeling the actual plasmids with multiple labels is that the redundancy increases the odds that a signal will be received from each molecule. Finally, the reliability of *in vitro* measurements of the theoretical parameters could be tested. The fractional occupation could alternatively be measured in *vivo* by working with a low copy plasmid that could be spatially resolved via conventional microscopy, then performing a photobleach experiment to determine the number of probes hybridized to a single plasmid.

We have also shown that, by increasing the number of labels on a target, one can take advantage of the central-limit theorem to improve on the accuracy of a molecular count, to achieve a 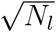 improvement in the uncertainty of the estimated count (where *N_l_* is the mean number of labels per molecule). This approach is well adapted for counting plasmids because standard techniques for detecting specific DNA sequences, such as DNA FISH, require labeling with many fluorophore conjugated probes. However, the example is rather specialized, and it’s often not feasible to attach multiple fluorophores to a single-molecule, such as when directly expressing fluorescently tagged proteins. On the other hand, standard immunolabeling techniques regularly target proteins with multiply labeled, fluorescently conjugated antibodies in order to achieve good signal intensity. Redundant labeling would reduce the uncertainty in quantifying the number of molecular components within diffraction limited clusters or aggregates via this commonly employed imaging technique.

## Appendix A: Negative binomial distribution

Identifying Eq. 2 as a negative binomial distribution, the mean and variance can easily be derived from the moment generating function:

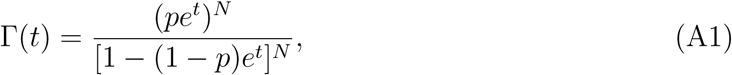
where *t* is a dummy variable and, for consistency with Eq. 2, *p* = 1 − e^−1/λ^. The *k^th^* moment is solved for by evaluating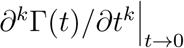. From the first two moments, we once again obtain Eqs. 3 and 4 for the mean and variance, respectively.

## Appendix B: Derivation of the mean number of plasmids

To derive equation 13, we begin by expressing *p*(*M*|B) analogous to Eq. 10 as

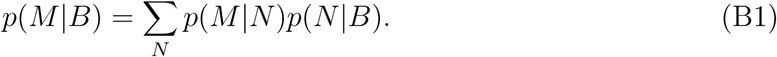

To calculate the mean, we can multiply both sides by ∑*_M_ M* such that

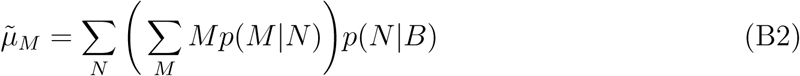

We approximate the term in brackets with an estimate of the expectation value of *p*(*M*|*N*), which is *N/(*θ*h*). This leaves

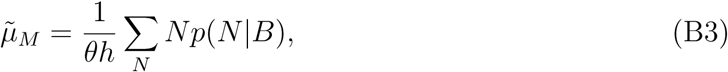
where the remaining sum is identified as *μ_N_*. Substituting the expression we derived in Eq. 6 yields Eq. 13.

## Appendix C: Accuracy of the estimate

If we Taylor expand the log likelihood function 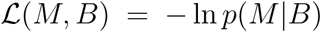, assuming the distribution to be peaked about 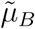, we can relate the log likelihood function to the variance in the measured number of blinks [28]:

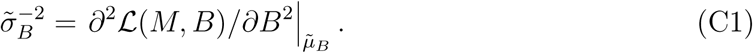

However, we are interested in calculating the variance in the number of molecules 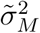, so let’s assume that the probability distribution *p*(*M*|*B*) is also peaked about 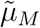, and approximate its functional dependence as a Gaussian centred at 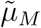 with variance 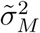 (i.e., Laplace approximation). In this case, the log likelihood function may be expressed as follows:

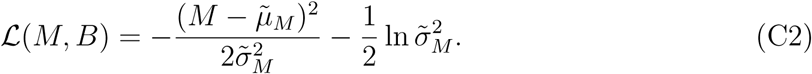

Since 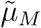 and 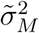 are both functions of *B*, we can evaluate the second derivative of Eq. C2 to obtain:

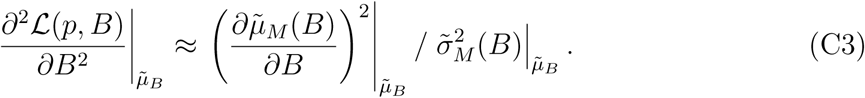

Combining this result with Eq. C1 and solving for 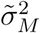 yields Eq. 14.

## ACKNOWLEDGMENTS

This work was funded by the Natural Sciences and Engineering Research Council of Canada [J.N.M., D.N., A.Z.], an Early Researcher Award from the Ministry of Research and Innovation [J.N.M., N.R.], the Human Frontier Science Program [Y.W.], and by the Arkansas Biosciences Institute, the major research component of the Arkansas Tobacco Settlement Proceeds Act of 2000 [Y.W.]

## AUTHOR CONTRIBUTIONS

D.A.N., Y.W. and J.N.M. conceived of the study. D.N., N.R. and J.N.M. performed the simulations. D.N., J.N.M., and A.Z. developed the analytical theory. All authors contributed to writing the manuscript.

